# Dose-dependent suppression of hippocampal contextual memory formation, place cells, and spatial engrams by the NMDAR antagonist (R)-CPP

**DOI:** 10.1101/2022.06.13.495957

**Authors:** Mengwen Zhu, Mark G. Perkins, Richard Lennertz, Alifayez Abdulzahir, Robert A. Pearce

## Abstract

A common way to study the functional importance of N-methyl-D-aspartate receptors (NMDARs) in hippocampal memory-encoding circuits is by administering NMDAR antagonists. We recently compared the effects of (R,S)-3-(2-carboxypiperazin-4-yl)-propyl-1-phosphonic acid (CPP), a competitive NMDAR antagonist, on suppression of memory in vivo versus suppression of NMDAR-mediated field EPSPs (fEPSP_NMDA_) and long-term potentiation (LTP) in vitro. Surprisingly, we found that concentrations that block contextual conditioning *in vivo* are ineffective at blocking the fEPSP_NMDA_ or LTP *in vitro*. Here we tested one possible explanation for the mismatch – that the hippocampus is relatively resistant to CPP compared to other brain structures engaged in contextual fear conditioning. We used the context pre-exposure facilitation effect (CPFE) paradigm to isolate the hippocampal component of contextual learning, and in-vivo calcium imaging of place cells and spatial engrams to directly assess hippocampal spatial coding. We found that, by both measures, the active enantiomer (R)-CPP did interfere with hippocampal function at concentrations below those that block fEPSPs or LTP. We conclude that the alternative – that CPP interferes with memory by targeting NMDARs in interneurons rather than pyramidal neurons – is the more likely explanation.

## 1. Introduction

N-methyl-D-aspartate receptors (NMDARs) are ionotropic glutamate receptors that mediate excitatory synaptic transmission throughout the central nervous system (CNS) (Paoletti et al., 2013). Multiple subtypes of this tetrameric receptor exist, composed of subunits that exhibit temporally and spatially dynamic expression patterns. NMDARs are essential for learning and memory and other higher cognitive functions. Accordingly, their dysfunction has been implicated in a number of neurological disorders, including Alzheimer’s disease, Huntington’s disease, schizophrenia, and neuropathic pain (Balsara et al., 2014; Chou et al., 2020; Kellermayer et al., 2018; Lau and Zukin, 2007; Paoletti, 2011; Zhou et al., 2011; Zhou et al., 2013).

Many NMDAR-selective agonists, antagonists, and allosteric modulators have been developed over the past several decades. They have been used to probe the functional roles of NMDARs in different cognitive processes and pathological conditions and tested for their potential therapeutic value (Ahmed et al., 2020). Studies using NMDAR antagonists in particular have revealed a wealth of information about how the molecular composition, spatial and temporal expression patterns, gating kinetics, and pharmacological sensitivities of various NMDAR subtypes influence their physiological roles (Feng et al., 2005; Hansen et al., 2014; Hansen et al., 2018; Lind et al., 2017; Paoletti et al., 2013; Stroebel et al., 2018; Wyllie et al., 2013; Zhu and Paoletti, 2015). One such antagonist, (R,S)-3-(2-carboxypiperazin-4-yl)-propyl-1-phosphonic acid (CPP), has been widely used for decades, primarily because it readily crosses the blood-brain barrier so it can be administered systemically, but also because it is water soluble, making it convenient to use both *in vitro* and *in vivo* (Bergeron and Rompre, 2013; Fung et al., 2016; Gemperline et al., 2014; Harris et al., 1986; Hayashi, 2019; Laha et al., 2022; Lehmann et al., 1987; Wang et al., 2021).

In a recent study, we sought to use CPP to establish the quantitative relationship between NMDAR block, interference with long-term potentiation (LTP) *in vitro*, and suppression of contextual memory *in vivo* (Laha et al., 2022). We found a good correspondence between concentrations of CPP required to block NMDAR field EPSPs (fEPSPs; IC50 = 434 nM) and suppress LTP in hippocampal brain slice preparations (IC50 = 361 nM). However, these concentrations were nearly an order of magnitude greater than the concentration of CPP (53 nM) corresponding to the dose that interferes with contextual fear conditioning *in vivo* (2.3 mg/kg).

In the present study we sought to test one possible explanation for this mismatch: that CPP blocks contextual fear conditioning not by interfering with hippocampal function, but by interfering with non-hippocampal processes, such as the amygdala-dependent association of the context with the aversive stimulus. We performed two types of experiments. First, we compared the concentrations of (R)-CPP required to block hippocampal versus non-hippocampal components using the context pre-exposure facilitation effect (CPFE) paradigm, a modified version of contextual fear conditioning that separates the hippocampus-dependent contextual learning from the amygdala-dependent association between the recalled context and the aversive shock. Second, we measured the effects of (R)-CPP on the formation and stability of hippocampal place cells and spatial engrams as neural correlates of contextual memory. We found that the hippocampus- and amygdala-dependent components of CPFE were equally sensitive to suppression by low doses of (R)-CPP, and that these doses also interfered with place cell and spatial engram formation. We conclude that (R)-CPP blocks memory formation by interfering with hippocampal function, but that it does so by modulating NMDARs at sites that are not engaged *in vitro* in the same manner that they are *in vivo* – perhaps through interneuron circuits that do not contribute to fEPSPs and are not required to elicit LTP using standard induction protocols *in vitro*, but are essential for successful mnemonic function *in vivo*.

## 2. Materials and methods

All studies were conducted under Institutional Animal Care and Use Committee (IACUC) approval in accordance with the National Institutes for Health (NIH) guide for the care and use of laboratory animals (NIH Publications No. 8023, revised 1978).

### 2.1. Context Pre-exposure Facilitation Effect (CPFE)

A total of 88 C57BL/6J mice (half of each gender; Jackson Laboratories, Bar Harbor, ME) in the age range of 70-80 days were used to study the effects of (R)-CPP on contextual learning and memory. Mice were housed four per cage (food and water available ad libitum) within the Waisman Center Rodent Models Core facility at the University of Wisconsin-Madison. The light-dark cycle was kept at 12:12-hr with lights turned off at 7 p.m. All experiments were performed during the light cycle. For one week prior to initiating the three-day CPFE experiment, mice were handled for two to three minutes per day in the behavioral testing room to reduce their anxiety, and on each experimental day, mice within their home cages were brought into the behavioral testing room to habituate for 30 min.

The CPFE learning paradigm was carried out over three days. On day 1, a mouse was placed in the testing chamber for a 10-minute context preexposure. The testing chamber (20 × 20 × 30cm) was constructed of clear acrylic with checkerboard-patterned paper covering three of the four walls and an electric-shock grid floor consisting of stainless-steel bars (diameter of 2mm) that were regularly spaced 1cm apart (Fig. 1A). On day 2, the mouse was placed back into the same testing chamber and after 15 sec was administered an aversive electric foot shock (2 sec, 1 mA). The mouse remained in the chamber for an additional 30 sec (total of 47 sec) and it was then returned to its home cage. On day 3, the mouse was again placed into the same testing chamber for 10 min and its freezing behavior was recorded via FreezeFrame™ software. The percentage of time it was ‘immobile’ on day 3 (% freezing) served as a measure of the strength of contextual fear memory.

**Fig. 1.**
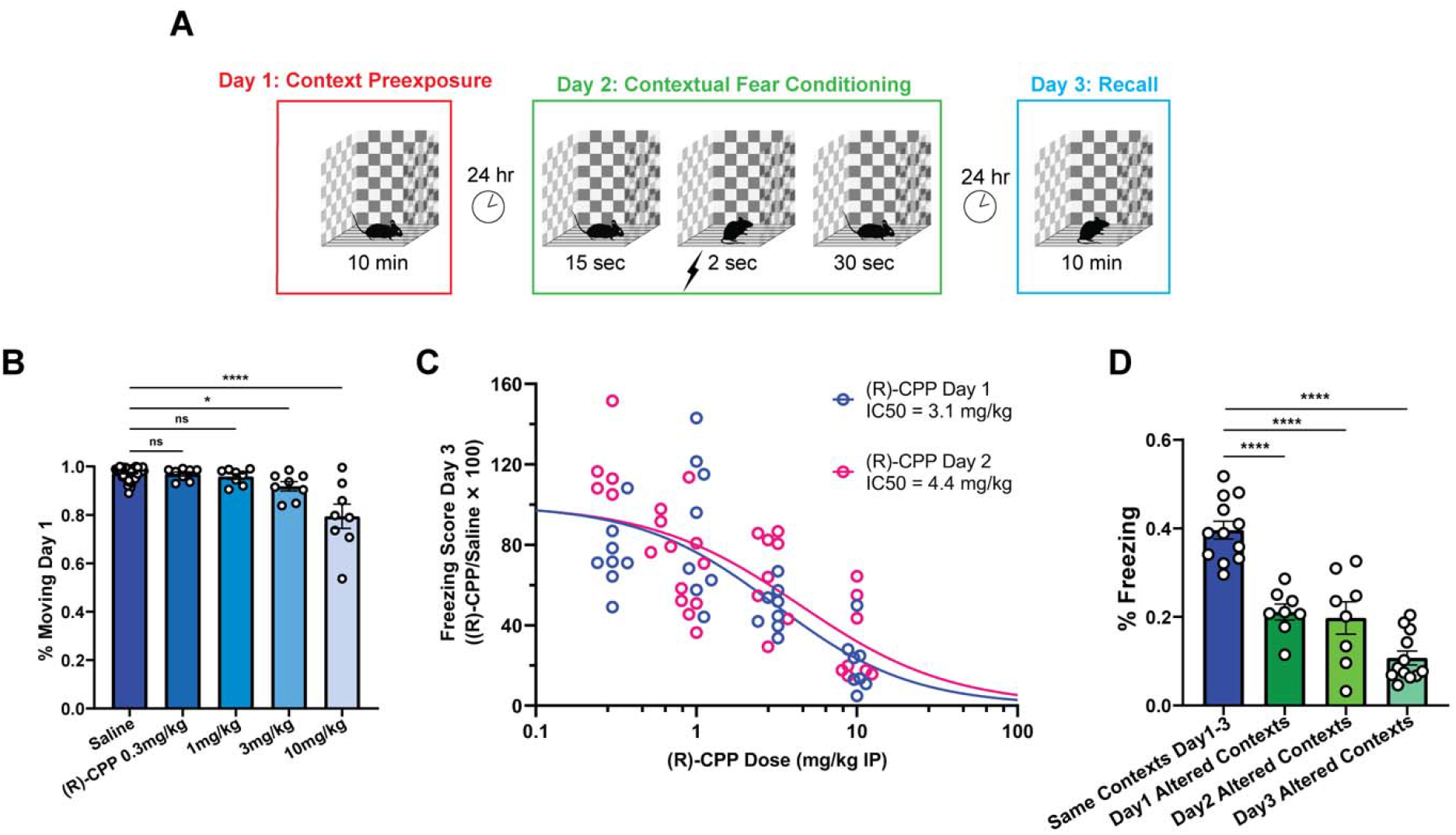
Dose-dependent suppression of contextual fear memory through (R)-CPP as measured by CPFE paradigm. A) Schematic depiction of CPFE paradigm. Pre-exposure groups were injected with saline or (R)-CPP an hour prior to day 1, and pre-shock groups are injected an hour prior to day 2. B) (R)-CPP dose-dependently suppresses mobility of the mice during novel context preexposure (day 1) (mean ± sem; each circle represents one mouse). C) Fitted (R)-CPP dose-response curves of both pre-exposure (blue) and pre-shock (magenta) groups in CPFE experiments. D) Presenting mice with altered contextual cues on day 1, day 2, or day 3 significantly reduces freezing response on day 3 compared to exposing mice consistently to the same context (mean ± sem).

To assess the effect of (R)-CPP on the ability of a mouse to form a memory of the context, mice were intraperitoneally (IP) injected with saline (control) or a range of doses (0.3, 1, 3, or 10mg/kg) of (R)-CPP (Tocris Bioscience, Bio-Techne Corp., Minneapolis, MN, USA) 60 min before context pre-exposure (day 1). To assess the effect of (R)-CPP on the ability of a mouse to associate the context with footshock, mice were intraperitoneally (IP) injected with saline (control) or a range of doses (0.3, 0.6, 1, 3, or 10mg/kg) of (R)-CPP 60 min before contextual recall and foot electric shock (day 2). Finally, to assess the specificity and generalization of contextual fear memory, we exposed additional groups of mice to different contexts (checkerboard wall was replaced by white non-transparent wall; electric-shock grid floor was replaced by white non-transparent plastic board) on day 1, day 2, or day 3 of the CPFE paradigm, and evaluated their freezing responses on day 3.

### 2.2. In-vivo calcium imaging

To assess the effects of (R)-CPP on hippocampal place cells and spatial memory traces (spatial engrams), we used a miniaturized fluorescent endoscope (nVoke, Inscopix Inc., Palo Alto, CA, USA) to measure calcium activity in the dorsal hippocampus of freely-exploring mice (Ghosh et al., 2011; Yang and Yuste, 2017) and simultaneously tracked the position of the mice as they explored a novel or familiar environment (Sheintuch et al., 2020; Ziv et al., 2013).

To prepare for *in vivo* calcium imaging recordings, four C57BL/6J mice underwent two stereotaxic surgeries separated by 2-3 weeks. For both procedures, mice were anesthetized with isoflurane ∼2% adjusted to maintain immobility and spontaneous ventilation, and warmed by a block heater to maintain normothermia. For the first surgery, a small craniotomy window was made over the right dorsal hippocampus (from bregma: AP =-2.0, ML =-1.6) and 500nL of virus carrying the genetically encoded calcium indicator GCaMP6f driven by the CaMKIIa promoter (Inscopix Ready-to-Image AAV1-CaMKIIa-GCaMP6f) was injected at DV =-1.6 at a rate of 80nL/min using a NanoFil syringe with a 35g needle driven by a UMP3 microsyringe pump and SMARTouch controller (WPI). The needle was kept in place for 2min, then retracted 200um, where it remained for an additional 8min before removing it slowly. The skin was sutured closed and the mouse was returned to its home cage. For the second surgery, a larger craniotomy was made at AP=-2.2, ML=-2.1, and the cortex corpus callosum overlaying the hippocampus was aspirated using a 30g blunt needle and cold saline irrigation, taking care not to aspirate the alveus. Bleeding was controlled with Gelfoam and cold saline irrigation. An integrated GRIN lens (1mm diameter x 4mm length) and baseplate were then inserted slowly at a 9-degree angle to the midline, until the center of the surface of the lens rested at DV=-1.2. The baseplate was cemented in place using Metabond, the surrounding skin was sutured closed, and the mouse was allowed to recover from surgery and anesthesia, then returned to its home cage. After each of the surgeries, mice received subcutaneous injections of 5mg/kg carprofen for pain control.

Two to three weeks after baseplate attachment, mice were screened for the quality of calcium fluorescence signals. The nVoke camera was affixed to the baseplate, and the mouse was placed into a recording arena (40 × 40 × 30cm acrylic enclosure) surrounded by blackout curtains and allowed to explore freely for 10min. A commutator (Inscopix Inc.) was used to prevent tangling of the wire attached to the microscope. During this initial ‘screening session’, the focal plane, LED power, and gain were adjusted within Inscopix Data Acquisition Software (IDAS 2019) to optimize recording parameters. This screening recording was processed with MATLAB API packages embedded within Inscopix Data Processing Software (IDPS v1.6.0; details described in section 2.3) to determine the number of cells with sufficient activity level (thresholds: >80 cells, each with >5 calcium events over 10min). Once sufficient activity was observed, the same settings were used for all subsequent recording sessions.

To measure the formation of place cells and spatial engrams, we conducted multiple pairs of 10-min recordings for each mouse, with the two sessions separated by 24 hours (referred to as session 1 and session 2). For session 1, the nVoke camera was attached and the mouse was placed in the recording arena used for screening sessions, but the arena included a unique set of visual, olfactory, and tactile cues. Visual cues consisted of 8 × 11’’ paper sheets with various colors on the outside of all four walls. Olfactory cues consisted of 1μL of odorant (benzaldehyde, hexanal, *a*-pinene, heptaldehyde, eugenol, or eucalyptol) on filter paper placed within covered culture dish in one corner of the arena. Tactile cues consisted of acrylic squares on the arena floor, various sizes of shallow glassware, and a sheet of exam table paper or hardware cloth covering the entire arena floor. To capture the position of the mouse, a bank of infrared lights (12 lux) was used to illuminate the arena so that the mouse could be tracked using an IR-sensitive camera (Basler aca1300-60gm) below the arena. The video camera and epifluorescent microscopy acquisition were synchronized using a hardware trigger controlled by EthoVision XT15 software (Noldus Information Technology). Immediately after the recording, the miniature microscope was detached, the mouse was returned to its home cage, and the arena was cleaned by replacing the exam table paper and wiping down all surfaces with EtOH. Session 2 took place 24 hours after session 1, using the exact same set of sensory cues as session 1. Each mouse underwent 5-6 pairs of recording sessions, with at least 48 hours between pairs of sessions.

To measure the effect of (R)-CPP on the formation of place cells and spatial engrams, 60 min before session 1 we injected IP saline (control) or the same doses of (R)-CPP (1, 3, and 10mg/kg) that had been used prior to context pre-exposure in CPFE experiments, and then carried out session 2 recordings without any injection. In a separate set of experiments, to measure the effect of (R)-CPP on the retrieval of spatial engrams, we omitted the session 1 injection and instead injected 10mg/kg (R)-CPP 60 min before session 2. To quantify the stability of spatial engrams, in some experiments we waited 72 rather than 24 hours before carrying out session 2 recordings. To establish specificity or generalization of spatial learning, for some experiments we presented a different set of sensory cues for session 2 (so-called ‘alternate contexts’ experiments).

### 2.3. Place cell and spatial engram analysis

Calcium imaging recordings were processed in two stages. The first stage utilized the MATLAB API package embedded within Inscopix Data Processing Software v1.6.0 (IDPS v1.6.0). Each recording was both spatially and temporally down-sampled by a factor of two and then ‘preprocessed’ to remove artifacts. Preprocessed recordings were spatially filtered (low cutoff = 0.005 pixel^-1^; high cutoff = 0.5 pixel^-1^) for better contrast and smoother frames, and then ‘motion corrected’ based on a region of interest (ROI) that contained blood vessels as background references. To better visualize calcium events and generate a maximum projected image for the estimation of average cell diameter (needed for subsequent analysis using the constrained nonnegative matrix factorization [CNMF-E] algorithm), a ‘ΔF/F’ transformation was applied so that fluorescent signals were quantified by deviations from background noise. The CNMF-E image analysis method was applied to the ΔF/F-transformed recording (Zhou et al., 2018), with analysis parameters tested and optimized within the IDPS v1.6.0. The noisy calcium traces of detected cells were deconvolved using the ‘online active set method to infer spikes’ (OASIS) method (Friedrich et al., 2017). Once deconvolved, calcium traces were event-detected based on biokinetics of GCaMP6f using a threshold of 4 median absolute deviations (MAD) to establish ‘cellsets’ of calcium events (Chen et al., 2013). Analyzed cellsets were randomly chosen to visually verify that inferred spikes accurately reflected calcium signals from detected cells. To be included in the second stage of analysis, only cells with exactly one spatial component, appropriate size, and more than 5 detected calcium events over 10 minutes of recording were accepted. Finally, cellsets of all recordings obtained from each individual mouse were ‘longitudinally registered’, so that the activity of each unique cell could be traced across all recording sessions (Sheintuch et al., 2017). These data were exported as csv files that included cell global ID, the timestamps (beginning of rising phase) of inferred calcium spikes, and the amplitude of calcium spikes in MAD.

For the second stage of analysis, we used custom-written MATLAB functions to merge behavioral tracking data and CNMF-E cellsets in order to characterize place cells and spatial engrams. ***Behavioral tracking analysis***: (1) speed was calculated based on mice’s XY-locations and sampling frequency of Noldus behavioral tracking system, and then smoothed with a triangular window (size = 500 msec); (2) the fraction of time that the mouse spent actively exploring (speed > 2 cm/sec) was calculated and reported as % moving; (3) an occupancy-time map (sec) was created by dividing the arena into a 10-by-10 matrix (i.e. 100 pixels), and computing the duration that a mouse spent within each pixel. ***Place cell analysis***: (1) calcium event timestamps were matched with behavioral tracking timestamps in order to find the spatial location of the mouse at the onset of each calcium event; (2) calcium events that occurred during periods of immobility (instantaneous speed < 2cm/sec) were excluded from analysis; (3) for each cell, an amplitude-weighted calcium event map was created by duplicating calcium events according to their amplitudes (∼1-8 MAD) and distributing them along the track of the mouse, at intervals of 40ms, over the rising phase of the GCaMP6f fluorescence signal; (4) for each cell, a *calcium event rate map* (in Hz; 10-by-10 matrix) was created by dividing the calcium event map by the occupancy-time map, pixel-by-pixel; (5) for each cell, the mutual information (MI) between the calcium event rate map and occupancy-time map for each cell was calculated according to the formulae:

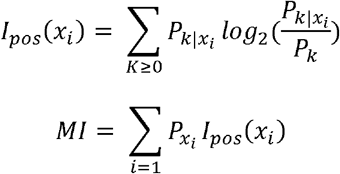

Where:

- *x*_*i*_ refers to the *i*^th^ pixel of the arena (in our analysis, the arena is binned into a 10-by-10 matrix).
- *I*_*pos*_ (*x*_*i*_) is the positional information (in bits) of pixel *x*_*i*_.
- 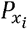 is the probability that the mouse occupies pixel *x*_*i*_, calculated as 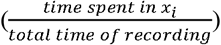.
- *P*_*k*_ is the probability of observing *k* calcium events
- 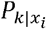 is the conditional probability of observing *k* calcium events in pixel *x*_*i*_.
- *MI* is the mutual information (in bits per second) between calcium event rate and mouse’s spatial location.

(adapted from: (Kinsky et al., 2018; Olypher et al., 2003)); (6) for each cell, a null distribution of time-shuffled MI values was created by 1000 random circular permutations of calcium event timestamps; this null distribution was used to calculate p(MI), which represented the fraction of shuffled MI values that fell above the actual MI value of the cell. ***Rate Map Correlation (RMcorr)***: (1) for each cell, calcium event rate maps were smoothed by a Gaussian filter (radius = 8 cm, sigma = 6 cm) to generate gaussian-smoothed calcium event rate maps; (2) for each cell, Pearson’s correlation coefficient (PCC) was calculated between the vectorized smoothed calcium event rate maps from any two recording sessions; (3) to detect and account for possible coherent rotation of cells’ event rate maps (Kinsky et al., 2018), we calculated RM correlation by rotating maps from one session by 0°, 90°, 180°, and 270°, and the rotation that resulted in maximum mean(RMcorr) was designated as the angle of coherent rotation—which usually was 0°; (4) for each pair of sessions, the distribution of PCCs from all coactive cells (i.e. those with >5 calcium events during both sessions) was referred to as RMcorr distribution; (5) for each pair of sessions, a null RMcorr distribution was created by calculating RMcorr between spatially-permutated pairs of smoothed calcium rate maps (each pair was shuffled 100 times). ***Population Vector Correlation (PVcorr)***: (1) for each pair of sessions, smoothed calcium event rate maps of all coactive cells were organized into two 3-dimensional matrices, thus forming for each pixel a pair of population vectors (PVs); (2) for each pixel, the PCC between the two PVs was calculated, resulting in a PV correlation distribution across all pixels; (3) to detect and account for coherent rotation, we calculated PV correlation by using the same rotation procedure as in RM correlation computation, and again an angle of coherent rotation based on PV is determined—which again usually was 0°; (4) for each pair of sessions, a null PVcorr distribution was created by calculating PVcorr between two randomly-permutated smoothed event rate map matrices (100 shuffles).

### 2.4. Statistical analysis

Statistical comparisons for CPFE studies were performed using one-way ANOVA followed by Dunnett’s multiple comparisons tests, and dose dependence was assessed by fitting data to a logistic equation (GraphPad Prism v9.3.1). Comparisons for calcium imaging data were made by fitting linear mixed effects models, with statistical significance derived from Kenward-Roger type III & likelihood ratio tests (Bates et al., 2015) using RStudio, 2021.09.0 Build 351 (R Core Team (2021). R: A language and environment for statistical computing. R Foundation for Statistical Computing, Vienna, Austria. URL https://www.R-project.org/.).

## 3. Results

### 3.1 Context Preexposure Facilitation Effect (CPFE)

To assess the effect of (R)-CPP on the hippocampal component of contextual fear memory formation, we injected mice IP with saline (control) or (R)-CPP (0.3, 1, 3, or 10 mg/kg) 60 minutes before they were pre-exposed to a novel experimental arena on day 1 of the CPFE paradigm (Fig. 1A). (R)-CPP produced a modest, but statistically significant (F(4, 91) = 26.03, p < 0.0001), reduction in exploratory activity on day 1 (Fig. 1B). The mice were reintroduced into the arena on day 2 and administered a foot shock after only 15 sec, and after 30 sec returned to their home cages. On day 3 they were again placed in the same arena, and the amount of time that the mice spent ‘freezing’ was taken as a measure of their ability to associate the arena with the aversive stimulus (Fig. 1C). (R)-CPP administered on day 1 produced a dose-dependent reduction in freezing on day 3 (blue circles and logistic curve; F(4, 39) = 21.84, p < 0.0001), with an IC50 of 3.1 mg/kg [95% CI 2.0-5.1 mg/kg]. This dose dependence is similar to that which we measured previously for suppression of standard contextual fear conditioning by CPP, which had an IC50 of 2.3 mg/kg (Laha et al., 2022).

Day 1 of the CPFE paradigm engages the hippocampus, but not the neural circuitry that transmits the aversive stimulus or the amygdala. To assess the sensitivity of these other components, we performed a second series of CPFE experiments in which we administered (R)-CPP (0.3, 0.6, 1, 3, or 10mg/kg) on day 2 (Fig. 1C, magenta circles). Again, (R)-CPP produced a dose-dependent suppression of % freezing on day 3 (magenta logistic curve; F(5, 42) = 22.20, p < 0.0001), with an IC50 of 4.4 mg/kg [95% CI 2.7-8.4 mg/kg] – essentially identical to the dose dependence when (R)-CPP was administered on day 1. Since (R)-CPP was present during the aversive conditioning phase in these experiments, the findings indicate that extra-hippocampal circuits do not display a greater sensitivity than hippocampus to suppression by (R)-CPP.

As controls, we performed a series of experiments similar to the CPFE experiments presented above, but with altered sets of contextual cues presented to mice on either day 1, day 2, or day 3 (Fig. 1D). In each case, the % freezing on day 3 was significantly reduced compared to mice that had been presented the same context every day (F(3, 36) = 34.73, p < 0.0001). These results indicate that the hippocampal representation of the context was specific to the environmental cues provided, that context pre-exposure overcame the immediate shock deficit (day 1 altered context), that fear memory was context-specific (day 2 altered context), and that contextual fear memory did not generalize (day 3 altered context).

### 3.2 In-vivo calcium imaging: place cells

The results presented above support a model in which (R)-CPP suppresses contextual memory by directly interfering with hippocampal function. To further test this model, and to gain additional mechanistic insight, we measured the effect of (R)-CPP on the hippocampal representation of the environment using a genetically encoded calcium sensor (GCaMP6f) and a head-mounted miniaturized fluorescent microscope (Inscopix nVoke) to monitor the activity of CA1 pyramidal neurons as mice freely explored a novel or a familiar environment. We simultaneously tracked the position of the mice within the arena using a video camera and behavioral analysis software (Noldus Ethovision XT).

We first examined the influence of (R)-CPP on the formation of place cells in a novel environment by IP injecting saline or (R)-CPP (1, 3, and 10 mg/kg) 60 minutes prior to novel context exposure (Fig. 2A), matching day 1 of the CPFE paradigm (Fig. 1A). Examples of the calcium signals observed in several representative cells over the course of a 10-minute recording are shown in Fig. 2B. As also seen in CPFE experiments (Fig. 1B), (R)-CPP caused a dose-dependent reduction in the mobility of mice (Fig. 2C, mobility ∼ drug + (1|mouse_ID), F(3, 13) = 12.6, p = 0.0004)), with significant effects at doses of 3 mg/kg (t(13) = -3.12, p = 0.0082) and 10 mg/kg (t(13) = -5.95, p < 0.0001). The overall calcium event rate remained unchanged (mean event rate ∼ drug + (1|mouse_ID) + (1|mouse_ID:date) + (1|cell_ID:mouse_ID), F(3, 12.68) = 2.95, p = 0.0734), but was reduced by 35.7% at a dose of 10mg/kg (t(11.68) = -2.71, p = 0.019), as shown in Fig. 2D.

**Fig. 2.**
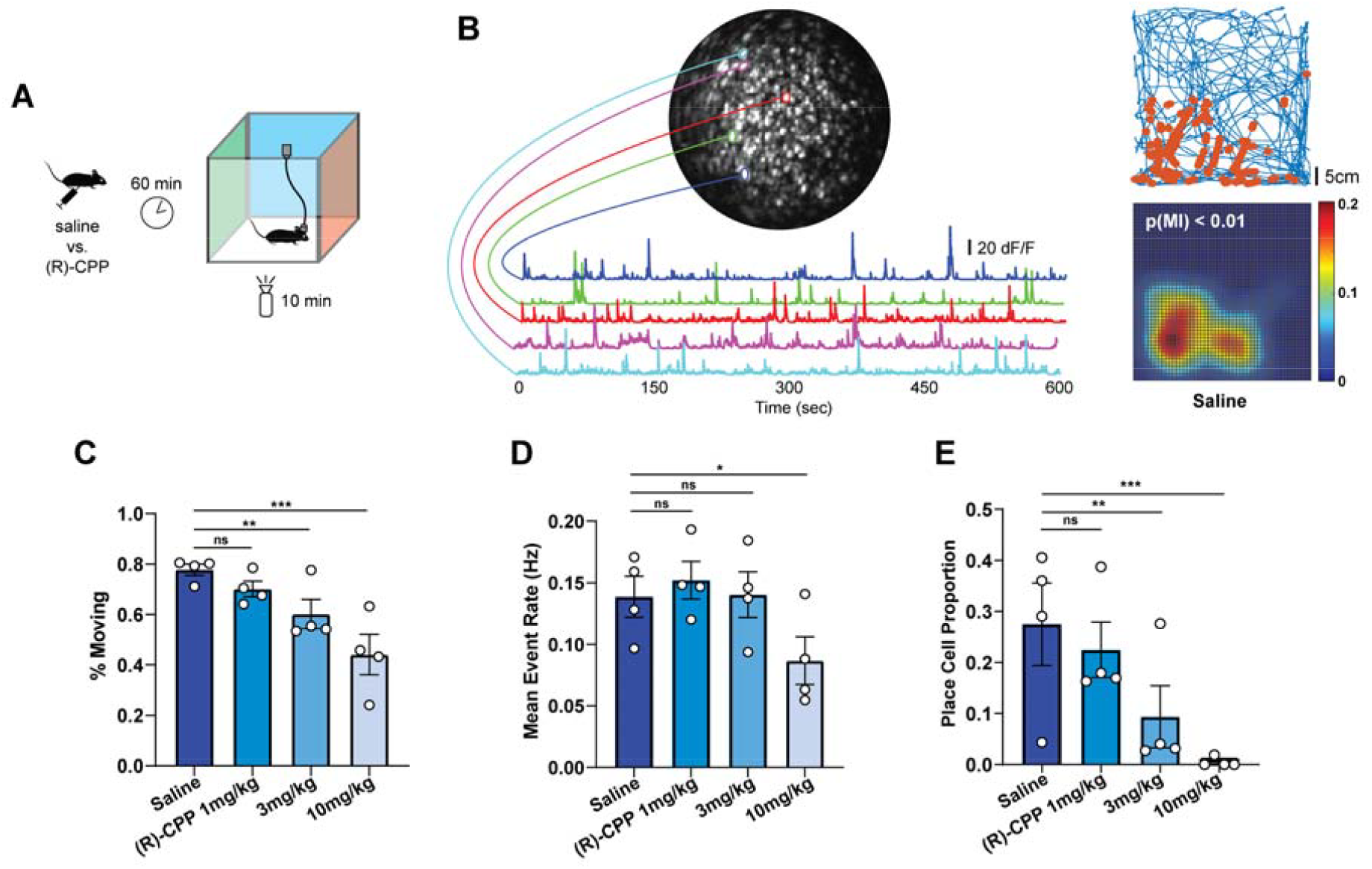
Dose-dependent suppression of place cell formation by (R)-CPP. A) Schematic depiction of the experimental paradigm used for measuring place cell formation in novel context exposure. Saline or (R)-CPP is administered an hour prior to context free exploration to match day 1 of CPFE paradigm. B) Maximum projected image of a 10-minute calcium imaging recording from one mouse and calcium traces of five selected cells (left). Example pairs of event dot map (top right) and gaussian-smoothed event rate map (lower right) of one detected cell when mouse is injected with saline (control). In event dot maps, the blue line represents the mouse’s motion trace and the strings of orange dots represents amplitude-adjusted calcium events. C) Dose-dependent reductions in mobility of the mice during free exploration that match the results from CPFE experiments (mean ± sem). D) Mean calcium event rates for all detected cells and all four mice (mean ± sem). Significant reduction was only observed at 10mg/kg (R)-CPP injection. E) Place cell formation, as defined by mutual information (MI) shuffle test, was dose-dependently suppressed by (R)-CPP in novel context exposure (mean ± sem).

To visualize the place-specificity of cell firing, for each cell we marked the location of the mouse when a calcium event was detected (Fig. 2B, red dots) over the track of the mouse as it explored the arena (blue line). From these data we derived a gaussian-smoothed event rate map for that cell (Fig. 2B, lower right panel). We then used the mutual information between the position of the mouse and the cell’s firing probability to determine whether the spatial modulation of firing rate was significantly greater than chance based on a permutation test. Under control conditions, 27±9 % of cells qualified as ‘place cells’. (R)-CPP reduced the proportion of place cells in a dose-dependent manner (Fig. 2E; % place cell ∼ drug + (1|mouse_ID) + (1|date), F(3, 16) = 11.75, p = 0.0003), with significant reductions seen at doses of 3 mg/kg (t(13) = -3.29, p = 0.0059) and 10 mg/kg (t(13) = -4.89, p = 0.0003). These results show that (R)-CPP interferes with hippocampal representation of space at the same doses that interfere with contextual memory formation, and suggests that (R)-CPP induces amnesia in contextual conditioning at least in part by suppressing place cell formation.

### 3.3 In-vivo calcium imaging: spatial engrams

Place cells are a well-recognized component of hippocampal spatial maps (Cobar et al., 2017; Eichenbaum, 2017; Howard and Eichenbaum, 2015). However, even cells that do not exhibit strong enough spatial modulation of firing to qualify as bona-fide place cells do exhibit context-specific firing rates and carry information about past exposure to a context (Tanaka et al., 2018), and their activity also contributes to the ‘spatial engram’ of the environment (Sheintuch et al., 2020; Stefanini et al., 2020). We therefore considered whether (R)-CPP might also interfere with development of stable spatial engrams.

To test the effect of (R)-CPP on spatial engram formation, we compared the spatially modulated activity of neurons during their initial exposure to a novel context (session 1, which were the recording sessions we analyzed for place cell development as described above) with their activity during re-exploration of that same context 24 hours later (session 2). This experimental design (Fig. 3A) thus mirrors the contextual learning on day 1 and contextual recall on day 2 of the CPFE experiment, after mice had been administered saline or (R)-CPP on day 1 (Fig. 1A). However, for these experiments no shock was administered, and the mice explored their environment for 10 min during recording session 2. Also, to measure the effects of (R)-CPP on spatial engrams, mice were tested repeatedly after different doses of (R)-CPP or saline before session 1, whereas in CPFE experiments each mouse underwent only a single 3-day experiment.

**Fig. 3.**
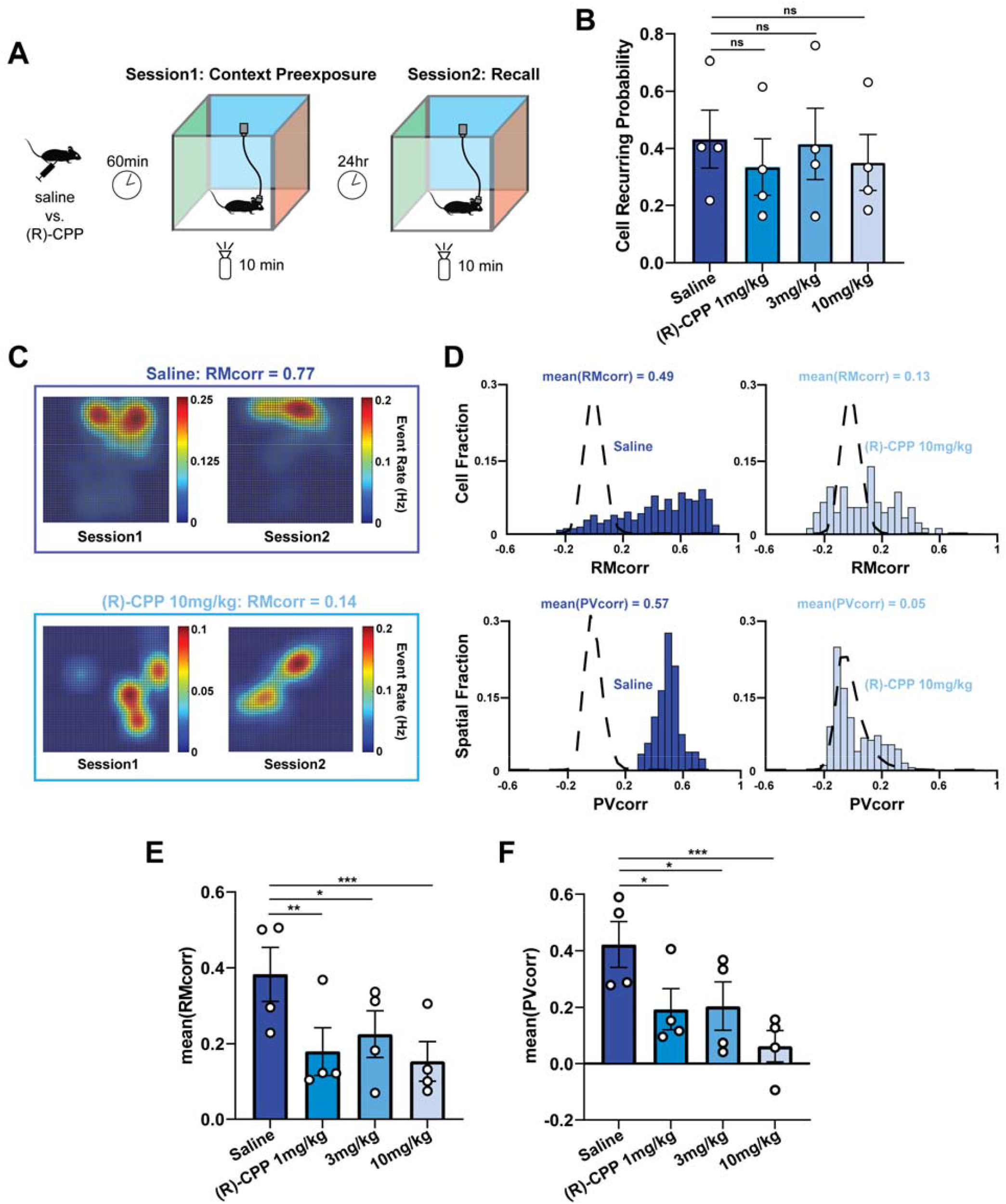
Dose-dependent suppression of spatial engram by (R)-CPP. A) Schematic depiction of the experimental paradigm used for evaluating spatial engram. Mice were injected with saline or (R)-CPP a hour prior to exposure to a novel context (session 1) and are re-exposed to the exact same context (session 2) to allow spatial engram retrieval. B) The probability that the same neurons became reactivated upon re-exposure to the same context was not influenced by (R)-CPP (mean ± sem). C) Example gaussian-smoothed rate maps formed on day 1 and day 2 of two cells in a mouse that is injected with saline (top) or 10mg/kg (R)-CPP (bottom). We observed more frequent ‘remapping’ of spatial firing patterns of neurons when mice are injected with (R)-CPP, as compared to saline baseline. D) Example RMcorr and PVcorr distributions from one mouse that is injected with saline (left) and 10mg/kg of (R)-CPP (right). The black broken line represents shuffled distributions (null). We observed striking shift in both RMcorr and PVcorr distributions toward null distribution when mice were injected with 10mg/kg of (R)-CPP. E) Dose-dependent reduction of mean RMcorr values when mice were injected with 1, 3, or 10mg/kg of (R)-CPP (mean ± sem). F) Dose-dependent reduction of mean PVcorr values when mice were injected with 1, 3, or 10mg/kg of (R)-CPP (mean ± sem).

As we also observed previously (Zhu et al., 2022), approximately one-half of cells that fired during session 1 also fired during session 2 (Fig. 3B). (R)-CPP did not alter the recurrence probability at any dose (cell recurring probability ∼ drug + (1|mouse_ID), F(3, 13) = 0.71, p = 0.5613; p > 0.1 for all doses). To quantify the strength of spatial engrams, we employed two computational strategies: rate-map correlation (RMcorr), which compares firing rate maps for individual cells on a cell-by-cell basis, and population-vector correlation (PVcorr), which compares firing rate vectors of all cells on a location-by-location basis (please refer to section 2.3 for more details). Examples of gaussian-smoothed rate maps for individual cells are shown in Fig. 3C, taken from a mouse that had been given saline (upper panel) or 10mg/kg of (R)-CPP (lower panel) prior to session 1. The similarity between rate maps after saline yielded a high RMcorr value (0.77), and their dissimilarity after (R)-CPP administration led to a low RMcorr value (0.14). To visualize the strength of the spatial engram at an ensemble level, distributions of all such RMcorr (and PVcorr) values were plotted and their means computed (Fig. 3D). In the example shown, the distributions of RMcorr and PVcorr values (Fig. 3D, left panels) following saline administration fell largely outside the null distributions (dashed lines) calculated by randomly shuffling firing event times, consistent with the formation of a stable spatial map during session 1 and its recall during session 2. By contrast, administration of 10mg/kg of (R)-CPP (right panels) led to RMcorr and PVcorr distributions that fell largely within the null distributions, indicating the absence of a stable map as correlations were closer to chance levels. Summaries of RMcorr and PVcorr data for all pairs of sessions for all four mice administered saline or a range of doses of (R)-CPP are shown in Fig. 3E & 3F. Mean levels of both RMcorr (RMcorr ∼ drug + (1|mouse_ID) + (1|mouse_ID:date) + (1|cell_ID:mouse_ID), F(6, 25) = 6.0072, p = 0.0005) and PVcorr (PVcorr ∼ drug + (1|mouse_ID) + (1|mouse_ID:date), F(6, 28) = 4.25, p.= 0.0036) were significantly reduced at all doses of (R)-CPP – interestingly including at 1 mg/kg, which did not significantly reduce place cell formation (Fig. 2E).

We performed additional control experiments to test for context specificity, engram stability, and the effects of (R)-CPP on recall as opposed to encoding. To test for context specificity, we altered the contextual cues between sessions A and B (Figs. 4A & 4B). Altering the context significantly reduced both RMcorr (t(17.52) = -2.28, p = 0.0353) and PVcorr (t(21.99) = -2.51, p = 0.0203), demonstrating that the arena-specific cues provided information that was encoded as neuronal representations of context. To test for engram stability, we compared RMcorr using 24 vs. 72 hr intervals between sessions A and B (Fig. 4C, “72h Recall”). We found that RMcorr was not different at 72 hr vs. 24 hr intervals (t(20.39) = -1.066, p = 0.2987). To test effects of (R)-CPP on recall, we administered 10 mg/kg of (R)-CPP 60 minutes before session 2 rather than session 1 (Fig. 4C, “pre-recall”). (R)-CPP did not significantly reduce RMcorr (t(22.38) = -2.045, p = 0.0528), which forms a contrast with the strong suppression of RMcorr when 10mg/kg of (R)-CPP was administered before session 1 (t(22.47) = -4.381, p = 0.0002).

**Fig. 4.**
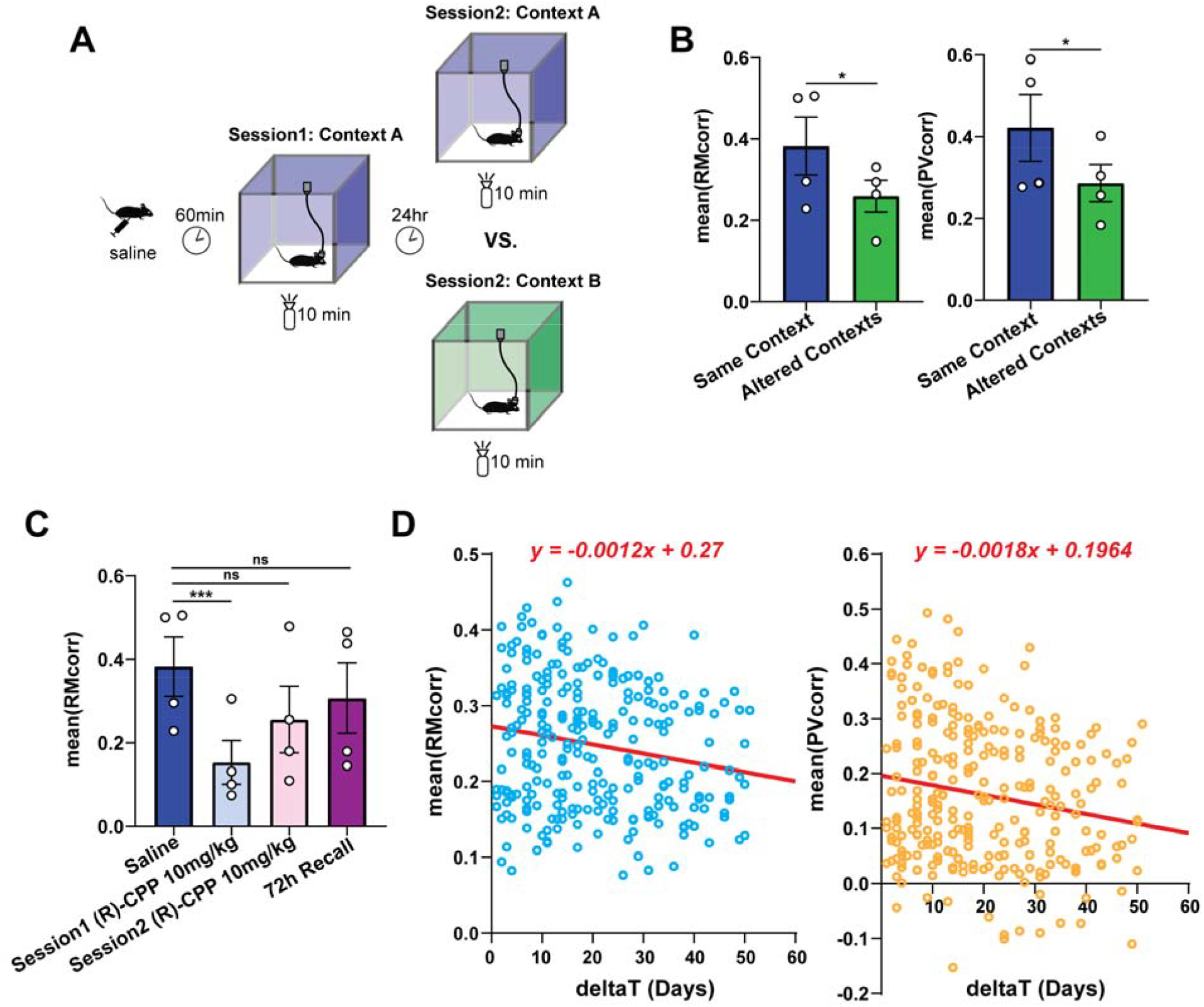
Altered contexts experiments and additional explorations on the properties of spatial engram. A) Schematic depiction of altered contexts experiments used to test whether spatial engram, as measured by RMcorr and PVcorr, was specific to the contextual cues we presented to the mice. B) Both RMcorr and PVcorr were significant reduced by altering contextual cues on day 2, indicating that spatial engram we measured was specific to local environment. C) A test on whether (R)-CPP also suppresses the recall of spatial engram (Pre-Day2 (R)-CPP 10mg/kg) and whether spatial engram lasts for more than 24 hours (72h Recall). 10mg/kg of (R)-CPP failed to block the retrieval of spatial engram, especially when compared to administering the same dose of (R)-CPP prior to day 1 (Pre-Day1 (R)-CPP 10mg/kg). Moreover, spatial engram remains stable beyond 24 hours (72h Recall). D) Exposing mice to novel contexts over a time span of ∼60 days reveals a significant drift in the residual correlation (RMcorr and PVcorr) of spatial engram. This may suggest a more general property of spatial engram that incorporates both local (changing) and distal (unchanging) contextual cues.

Athough 10mg/kg (R)-CPP induced complete amnesia as measured behaviorally via CPFE paradigm (Fig. 1D), we still observed a residual correlation that often fell outside of shuffled null distributions (Fig. 3D, E, & F). Furthermore, exposing mice to different contexts also did not completely eliminate RMcorr or PVcorr (Fig. 4B). These findings suggest that the spatial engrams we measured might incorporate both local (altered) contextual cues and more distal (relatively unaltered) contextual cues (e.g. experimenter, the experimental room, black curtains surrounding the arena, etc.). To test this hypothesis, and to evaluate how spatial engrams of distal contextual cues evolve over time, we examined RMcorr and PVcorr between pairs of recording sessions for mice that had been repeatedly exposed to novel contexts over time periods of up to 60 days (Fig. 4D). We found both values were lower for sessions that were separated by longer periods of time; fitting these data to a simple linear regression yielded significantly non-zero slopes (F(1, 266) = 9.761, p = 0.002 for RMcorr; F(1, 266) = 9.034, p = 0.003 for PVcorr). Therefore, spatial engrams do appear to include distal as well as proximal cues, and to drift to lower values over time.

## 4. Discussion

The present study was motivated by our recent finding that there is a substantial mismatch between the relatively high concentrations of CPP required to block fEPSPs (434 nM) and LTP (361 nM) versus the low concentrations that block contextual fear conditioning (53 nM) (Laha et al., 2022). Here we focused on one possible explanation – that CPP blocks contextual fear conditioning not by interfering with hippocampal function, but by interfering with some other essential process, such as the amygdala-dependent association of the context with the aversive stimulus. Our results indicate that this explanation is incorrect. In behavioral experiments that separated the hippocampus-dependent contextual learning phase from the amygdala-dependent association of context with the aversive stimulus, the dose-dependencies of contextual fear memory suppression were the same (Fig. 1C). Moreover, both matched the low doses of CPP that blocked standard contextual fear conditioning in our prior studies (Laha et al., 2022). The conclusion that low doses of (R)-CPP can interfere with hippocampal function specifically was reinforced by our calcium imaging results, which showed that (R)-CPP suppresses place cell formation with a dose dependence similar to its interference with contextual learning (Fig. 2E, c.f. Fig. 1C), and that these low doses also significantly reduce the expression of spatial engrams (Fig. 3E & F). Taken together, these results indicate that hippocampal function is indeed impaired at doses below those that block fEPSPs and LTP.

### 4.1 Non-hippocampal brain regions are not more sensitive than hippocampus to (R)-CPP

The CPFE paradigm relies on the concept of ‘immediate shock deficit’, wherein animals that are shocked immediately (within seconds) upon entry into a novel environment do not freeze on subsequent re-exposure, whereas mice that had been exposed on a prior day do exhibit a freezing response (Cushman et al., 2012; Fanselow, 2000; Miller et al., 2020; Rudy, 2009; Yavas et al., 2021). The deficit reflects the time it takes (several minutes) for the hippocampus to form a complete and retrievable contextual representation of an environment. However, once formed, it can be recalled quickly and associated with the aversive conditioning stimulus (Fanselow, 1990; Krasne et al., 2015).

We took advantage of this temporal separation between spatial learning and association with the aversive stimulus to compare the dose-dependent suppression of contextual memory formation by (R)-CPP (administration on CPFE day 1) versus contextual memory retrieval and context-fear association (R)-CPP (administration on CPFE day 2). The similarity in their dose-dependence (Fig. 1C) provides a clear indication that neither the aversiveness of the foot shock nor the amygdala-dependent association (day 2) exhibits a greater sensitivity to (R)-CPP than the hippocampus-dependent contextual learning (day 1).

### 4.2 Involvement of NMDARs in hippocampus-dependent contextual memory recall

Although the CPFE experimental design did allow us to conclude that the non-hippocampal components are not <u>more</u> sensitive than the hippocampus to (R)-CPP, we are unable to determine how sensitive they are, because the hippocampus is also engaged during the aversive conditioning phase (day 2) when the context is rapidly recalled. The essential involvement of the hippocampus in both the initial acquisition of contextual memory and subsequent memory retrieval was demonstrated in a prior study that employed muscimol, a GABA_A_R agonist, injected directly into the dorsal hippocampus prior to each of the three stages of the CPFE paradigm (Matus-Amat et al., 2004). That study did not address whether NMDARs specifically are required, but the findings that 10 mg/kg CPP did not suppress the reactivation of established place cells when rats were re-exposed to a familiar environment (Kentros et al., 1998), and that intrahippocampal infusion of D-APV did not impair spatial memory retrieval in a delayed matching-to-place paradigm (Steele and Morris, 1999), suggest that memory retrieval on CPFE day 2 might not require activation of NMDARs. However, other studies have shown that memory retrieval requires NMDAR activity-mediated AMPAR trafficking (Lopez et al., 2015), and that the excitability of engram cells at the moment of memory recall correlates with the success of context recognition (Pignatelli et al., 2019). Our present finding that there was a strong trend toward reduced RMcorr when 10 mg/kg (R)-CPP was administered before session 2 (recall) in calcium imaging experiments (Fig. 4C), further suggests that (R)-CPP might interfere with hippocampus-dependent rapid recall. In this case, the concentration-dependence for session 2 administration could reflect drug action on any of a number of brain structures, making it difficult to assign learning impairment to any one region or process specifically.

### 4.3 Place cells and spatial engrams as neural correlates of hippocampal memory

Recent years have seen tremendous advances in finding, tagging, and manipulating memory engrams – neuronal ensembles characterized by the expression of immediate early genes (IEGs) at the onset of active learning. These studies have provided compelling evidence that such ensembles are necessary and sufficient for the formation and retrieval of contextual memory (Asok et al., 2019; Chen et al., 2019; Eichenbaum, 2016; Ghandour et al., 2019; Hainmueller and Bartos, 2018; Liu et al., 2012; Tanaka et al., 2014; Tonegawa et al., 2015). However, these IEG-based methods do not provide information about the spatiotemporal dynamics of neuronal ensembles (Eichenbaum, 2016). Studies of place cells and spatial engrams, by contrast, can provide insight into this aspect of coding within the hippocampal memory-encoding circuit.

Place cells comprise a relatively small proportion of hippocampal pyramidal neurons that display spatially modulated firing patterns during free exploration (Cobar et al., 2017; O’Keefe, 1979). These spatial firing patterns, termed ‘place fields’, exhibit both stability and plasticity over long-time scales (e.g. days, weeks, or even months), thereby supporting context recognition, discrimination, and navigation (Gonzalez et al., 2019; Goode et al., 2020; Howard and Eichenbaum, 2015; Ziv et al., 2013). Place cells are thought to serve as a means by which the hippocampus encodes a navigable and annotatable ‘cognitive map’ (Eichenbaum, 2017; Moser et al., 2008). The ability to form and maintain an internal representation of the external world is proposed to serve as the basis for retrospective (e.g. memory recall) and prospective (e.g. future planning and simulation) cognition, and to render the hippocampus essential for the formation, consolidation, and retrieval of contextual memory (Lisman et al., 2017).

Evidence accumulated over several decades, including genetic and pharmacological manipulations that interfere with place cells and their long-term stability (Bannerman et al., 1995; Hayashi, 2019; Holahan et al., 2005; McDonald et al., 2005; Steele and Morris, 1999), indicates that NMDARs are important for this ‘cognitive map’. However, little has been done to quantitatively relate behaviorally relevant amnesia, or the disturbance of spatial information encoded within large neuronal ensembles, to the dose dependence of NMDAR blockade. Our present results demonstrating a clear dose dependence of place cell suppression by (R)-CPP (Fig. 2E), which matches the dose dependence of memory suppression in CPFE experiments (Fig. 1C), supports the concept that place cells are part of a cognitive map that undergirds memory, and that impairment of place cell formation by (R)-CPP contributes to its ability to suppress behaviorally relevant memory.

Like place cells, spatial engrams, as quantified by RMcorr and PVcorr (Zhu et al., 2022), are also defined by spatial modulation of firing rates. However, spatial engrams as defined in this manner include all cells that are active in both environments, not just place cells, because both have been shown to code for meaningful spatial information (Sheintuch et al., 2020; Stefanini et al., 2020; Tanaka et al., 2018). Also like place cells, spatial engrams can remain stable over periods of weeks (Alexander et al., 2020; Kinsky et al., 2018; O’Keefe, 1979; Olypher et al., 2003; Payne et al., 2021; Radvansky et al., 2021; Robinson et al., 2020; Sheintuch et al., 2020; Stefanini et al., 2020; Tanaka et al., 2018; Ziv et al., 2013). As ensemble spatial codes, spatial engrams fulfill some of the criteria that would qualify them as neural correlates of contextual memory. Our present results showing that spatial engrams are also suppressed by amnestic doses of (R)-CPP (Fig. 3) further supports their role in memory and suggests that suppression of spatial engrams also contributes to (R)-CPP-induced amnesia.

### 4.4 Hippocampal spatial engrams incorporate dissociable local and distal components and maintain long-term stability across days

The observation of residual RMcorr and PVcorr in altered contexts experiments (Fig. 4D) suggests that in addition to coding information about local contextual cues (altered), spatial engrams also code for distal contextual cues (unaltered). We were able to dissociate these two components of spatial engrams by exposing animals to novel local contexts (altered arena but unaltered room) over a span of ∼60 days and we observed significant drift in the residual RMcorr and PVcorr. This time-dependent slow evolution of ensemble spatial coding could possibly be explained by the spontaneously dynamic nature of place-cell activity (Hayashi, 2019). It may also relate to the process of memory consolidation that occur between hippocampus and the neocortex. According to systems consolidation theories, hippocampus acts as an initial store of engrams, which will be gradually transformed into more permanent memory traces within the neocortex upon repeated learning (Klinzing et al., 2019). Hence, one possibility that explains the drift in residual RMcorr and PVcorr is the gradual transformation of hippocampal representation of the room to a neocortical representation. Additionally, we report that the hippocampal representation of a two-dimensional arena remain highly stable at least within 72 hours, agreeing with previous studies using one-dimensional linear tracks (Sheintuch et al., 2020; Ziv et al., 2013).

### 4.5 Hippocampal interneurons may be targeted by (R)-CPP to produce behaviorally-relevant amnesia

Pharmacological suppression of hippocampal learning and memory through NMDAR antagonists has presumably been related to the suppression of LTP. However, our previous study demonstrated that there is a substantial mismatch in concentrations that block of NMDAR-dependent LTP in vitro and suppression of hippocampus-dependent contextual memory (Laha et al., 2022). Here, using CPFE, and supported by calcium imaging experiments, we established that the IC50 for (R)-CPP is 3.1 mg/kg (Fig. 1) and this dose caused significant disruption of hippocampal spatial coding (Fig. 2 & 3). Though we used (R)-CPP instead of CPP in the present study, the estimated free concentration of (R)-CPP in the brain is still much too low compared to the IC50 required to block LTP and fEPSP_NMDA_ in brain slices, taking into account the relative potency of (R)- and (S)-CPP and assuming that (R)-CPP has similar partition coefficient as its racemate (Aebischer et al., 1989; Feng et al., 2005; Gemperline et al., 2014).

Since our present results appear to rule out the possibility that a greater sensitivity of other brain structures can account for the mismatch, this leaves the actions of (R)-CPP on GluN2A-expressing interneurons as the most likely explanation (Zhu et al., 2022). GABAergic interneurons are key to maintaining the balance of synaptic inhibition and excitation within hippocampal memory-encoding circuits, and they themselves display diverse and subtype-specific expression of NMDARs (Booker et al., 2021; Huntley et al., 2020; Perszyk et al., 2016). In support of this possible explanation, it was reported recently that in hippocampal CA1 region, GluN2A subunits, which are preferentially depressed by (R)-CPP, are expressed in parvalbumin-positive (PV+) and somatostatin-positive (Sst+) interneurons and contribute to EPSCs at their glutamatergic synapses (Booker et al., 2021). Other studies also demonstrated the importance of interneuronal NMDARs in controlling temporal properties of dendritic feedforward inhibition and their involvement in malfunctioning brain circuits (Chittajallu et al., 2017; Maccaferri and Dingledine, 2002), but the subunit composition of their NMDARs is unknown. Future studies will be needed to directly test the role of NMDARs in specific types of interneurons in (R)-CPP-induced amnesia.

